# Hepatitis Viruses affect the expression of Endogenous Retrovirus and Tumor Microenvironment in HCC

**DOI:** 10.1101/2022.06.19.496748

**Authors:** Wei Zhang, Xiaoyang Wan

## Abstract

Hepatocellular carcinoma (HCC) is a commonly diagnosed cancer with high mortality rates.Chronic hepatitis B virus (HBV) infection is a major cause of hepatocellular carcinoma (HCC) world-wide. The molecular mechanisms of viral hepatocarcinogenesis are still partially understood. The immune response plays an important role in the progression of HCC. Immunotherapies are becoming an increasingly promising tool for treating cancers. Advancements in scRNA-seq (single-cell RNA sequencing) have allowed us to identify new subsets in the immune microenvironment of HCC. Yet, distribution of these new cell types and their potential prognostic value in bulk samples from large cohorts remained unclear. Here, we use single-cell RNA sequencing analysis to delineate the immune landscape and tumor heterogeneity in a cohort of patients with hepatitis viruses(HBV,HCV,HDV) associated human hepatocellular carcinoma (HCC).We also re-analyze the bulk RNA-seq data for the cohort to include the expression values of human Endogenous Retrovirus (hERV). And found correlations with hepatitis viruses status.

## Introduction

Liver cancer is the third leading cause of cancer-related mortality in the world (1), and hepatocellular carcinoma (HCC) accounts for approximately 90 percent of the incidence of all liver cancers. The niche of liver tissue includes various cell lines such as hepatocytes, endothelial cells, fibroblasts, epithelial cells, and immune cells. In this unique ecosystem, tumor formation requires coordinated functions and crosstalk between specific cell types. Compared with other types of cancer, the HCC-compliant tumor microenvironment has a strong dependence on the number and state of immune cells, resulting in a lack of clinical success in the treatment of HCC. Recent advances in this area have demonstrated that many factors are related to the clinical response to checkpoint therapy. Unfortunately, these results are implicit and have not been further developed as clinical biomarkers. A detailed understanding of the various immune cells in the tumor microenvironment (TME) is essential for the development of effective immunotherapy for HCC and the identification of new biomarkers. The immune microenvironment of HCC is a mixture of heterogeneous immune components, immune cells migrate from the hematopoietic organ to the liver, establish an active immune niche that interacts with stromal cells, and affect differentiation, tumorigenesis and development. Therefore, it is important to explore the composition and state of immune cells during tumorigenesis. T cells and B cells are the most abundant and most characteristic cell groups in solid tumor TME. B cells are always important cellular components in the tumor microenvironment, and play a role through antibody production, antigen presentation and immune regulation. The fundamental understanding of the subtle cellular and molecular landscapes in HCC remains elusive. Traditional strategies perform investigation mainly at the bulk-tumor level and have inherent limitations in providing precise information on individual cells residing in a highly admixed TME. Single-cell sequencing provides an important platform to study cancers. HCC is highly heterogeneous, causing current curative effect restricted (9). The bulk omics analysis technique is to sequence the tissue blocks, and for the cell types with low richness, the complexity of cells and intracellular changes will be obscured.

Single cell sequencing technology (such as single cell RNA-seq, scRNA) has been applied in liver cancer. In this study, we used cutting-edge algorithm provided by cellTypist [4] to re-analyze a public single-cell RNA sequencing dataset. CellTypist identified much more cell subtype than the default method. We also introduced human Endogenous Retrovirus [6] annotation file, which allow us to detect the expression level of human Endogenous Retrovirus from normal bulk RNA sequencing data. We identified some human Endogenous Retrovirus significantly differentially expressed among hepatocellular carcinoma with different hepatitis viruses status. We also found that the hepatitis viruses status are co-related with immune microenvironment in hepatocellular carcinoma.

## Materials and Methods

### 0.1 Single-cell transcriptomic analysis

Raw gene expression levels and clinical metadata for single-cell RNA sequencing (scRNA-seq) measurements from patients with Primary or Metastatic Hepatocellular Carcinoma were downloaded via Gene Expression Omnibus, through accession number GSE149614. The quality control was done using python package scanpy. [10] Cells annotation was conducted by python package CellTypist. [4]

### 0.2 bulk cell transcriptomic analysis

Raw fastq files (listed above) were downloaded from the GEO website. FastQC [13] v0.11.9 was used for quality control of the sequencing data, and Trimmomatic [2] v0.36 was used to trim out low quality reads. The gtf file contains 60 HERV families, across 14,968 loci was downloaded from RepeatMasker, combined with Comprehensive gene annotation file from GENCODE. The combined gtf file was used as annotation for the bulk RNA-seq data analysis. Reads were mapped to hg38 human genome using STAR [5] v2.7.7, and the abundance of genes was calculated using HTSeq2 [1] v0.13.5. DESeq2 [9] R package was used to identify differentially expressed (DE) genes.

### 0.3 Cell Deconvolution

Cell Deconvolution was conducted using R package MuSiC. [12] The annotated single-cell RNA-seq data mention above served as reference.

### 0.4 GSEA

GSEA GO enrichment analysis was conducted using R package clusterProfiler. [15]

### 0.5 Visualization

Volcano plot was generated using EnhancedVolcano R package. GSEA visualization was using enrichplot R package.

## Results

We re-analyzed the single-cell RNA-seq profiles from a total of ¿ 70,000 single-cell transcriptomes for 10 HCC patients [8]. We utilized the recently developed comprehensive reference of immune cell types program cellTypist (figure 1B) and annotated the 49386 cells in the study. The annotated cell types are much more than the leiden method in the Scanpy [14] or Seurat [7] package.

**Figure 1:**
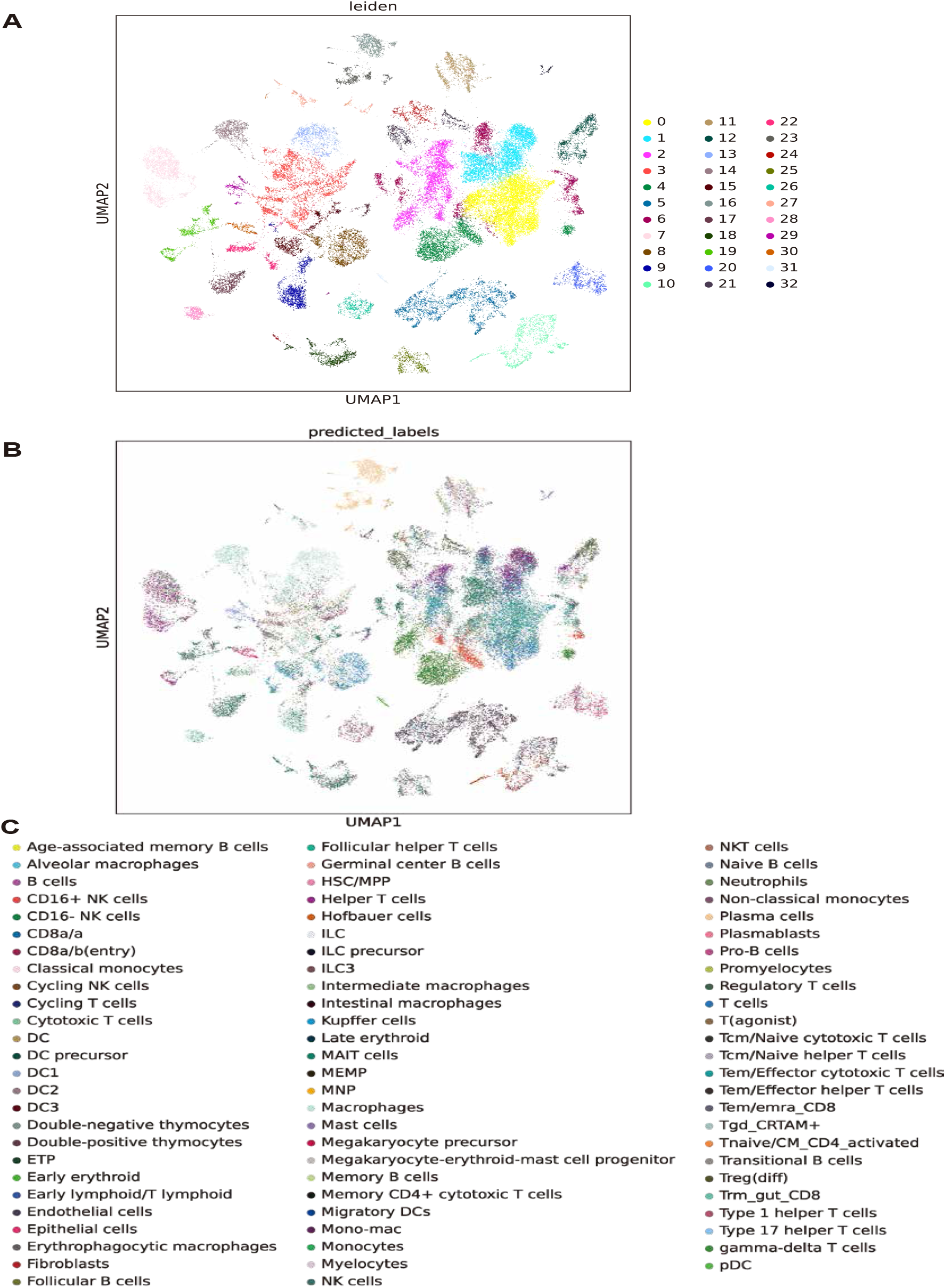
Re-analysis of scRNA-seq using cellTypist.

Mongolia has the highest incidence of hepatocellular carcinoma (HCC) in the world, but its causative factors and underlying mechanism remain unknown. We downloaded transcriptome sequencing of HCC from 76 Mongolian patients from GEO [3]. We used newly developed bulk RNA-seq deconvolves MuSiC [12] to obtain the proportions of these cell types in each sample. MuSiC incorporates “marker gene consistency” both cross-subject and cross-cell, to guard against bias in subject selection and cell capture in scRNA-seq. By incorporating both types of consistency, MuSiC allows for scRNA-seq datasets to serve as effective references for independent bulk RNA-seq datasets involving different individuals. The Figure2 showed the cell types proportions in each tumor samples, as well as other clinical characters including HBV,HCV,HDV infection status, with or without family history of liver cancer, smoke and alcohol habit, and the TNM stage information.

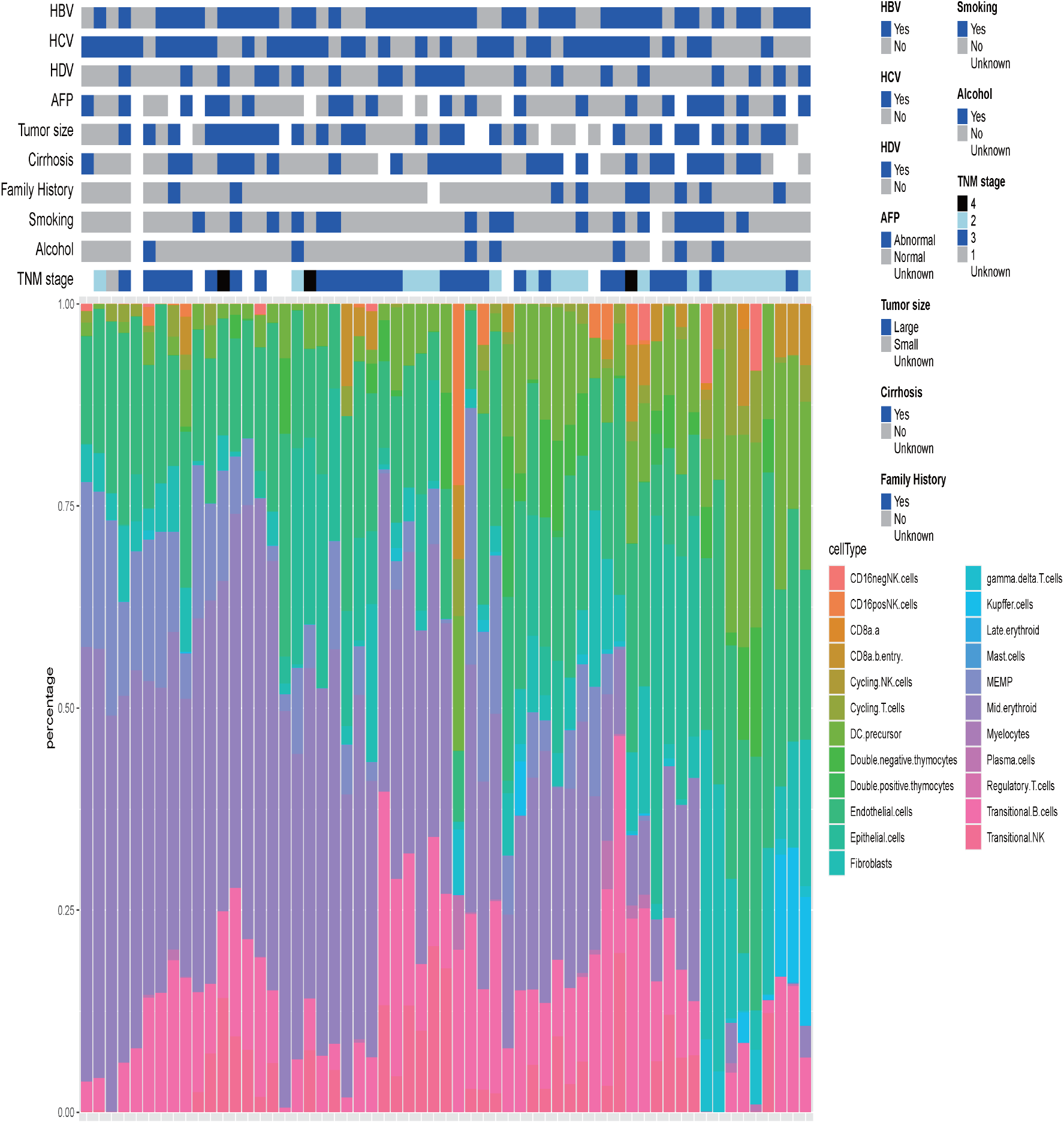

About 8 percent of human genome are come from retroviruses. hERVs are relics of ancient multiple infections that affected human germ line along the last 100 million of years, and became stable elements at the genome. Most hERVs are not active and functional due to the interruptions caused by multiple insertions/mutations. To investigate the expression of hERVs from RNA-seq data may find a missing piece to the human transcriptome jigsaw puzzle. Transcriptional units for HERV proviruses were defined by combining RepeatMasker annotations belonging to the same HERV subfamily that are located in adjacent or nearby genomic regions. Briefly, repeat families belonging to the same HERV subfamily (internal region plus flanking LTRs) were identified using the RepBase database. [11] We downloaded the human Endogenous Retrovirus annotations file from the RepBase database, and we combined the annotation file with ENCODE human annotation GTF file. Then we use the merged GTF file and human genome hg38 fasta file to build index for STAR aligner. [5] Then we used STAR and HTseq2 [1] to get the read counts for both genes and human Endogenous Retrovirus. We found that many human Endogenous Retrovirus are differentially expressed in tumor samples with different hepatitis virus status. For example, if we ingore the status of HCV and HDV, we can see that there are many differentially expressed genes and human Endogenous Retrovirus such as AFP, GABRG3, PGC, FREM2, HERV9.12p12.3 and HERV9.10p11.21 are significantly highly expressed in HBV positive hepatocellular carcinoma; LAMC2 and HARLEQUIN.10q22.1 are significantly lowly expressed in HBV positive hepatocellular carcinoma. We also did the GSEA using clusterProfiler. [15] All of the top six molecular pathway of the GSEA result are immune response related, Fig 3B. And all of the six molecular pathways are downregulated in HBV positive hepatocellular carcinoma comparing to HBV negative hepatocellular carcinoma.

**Figure 3:**
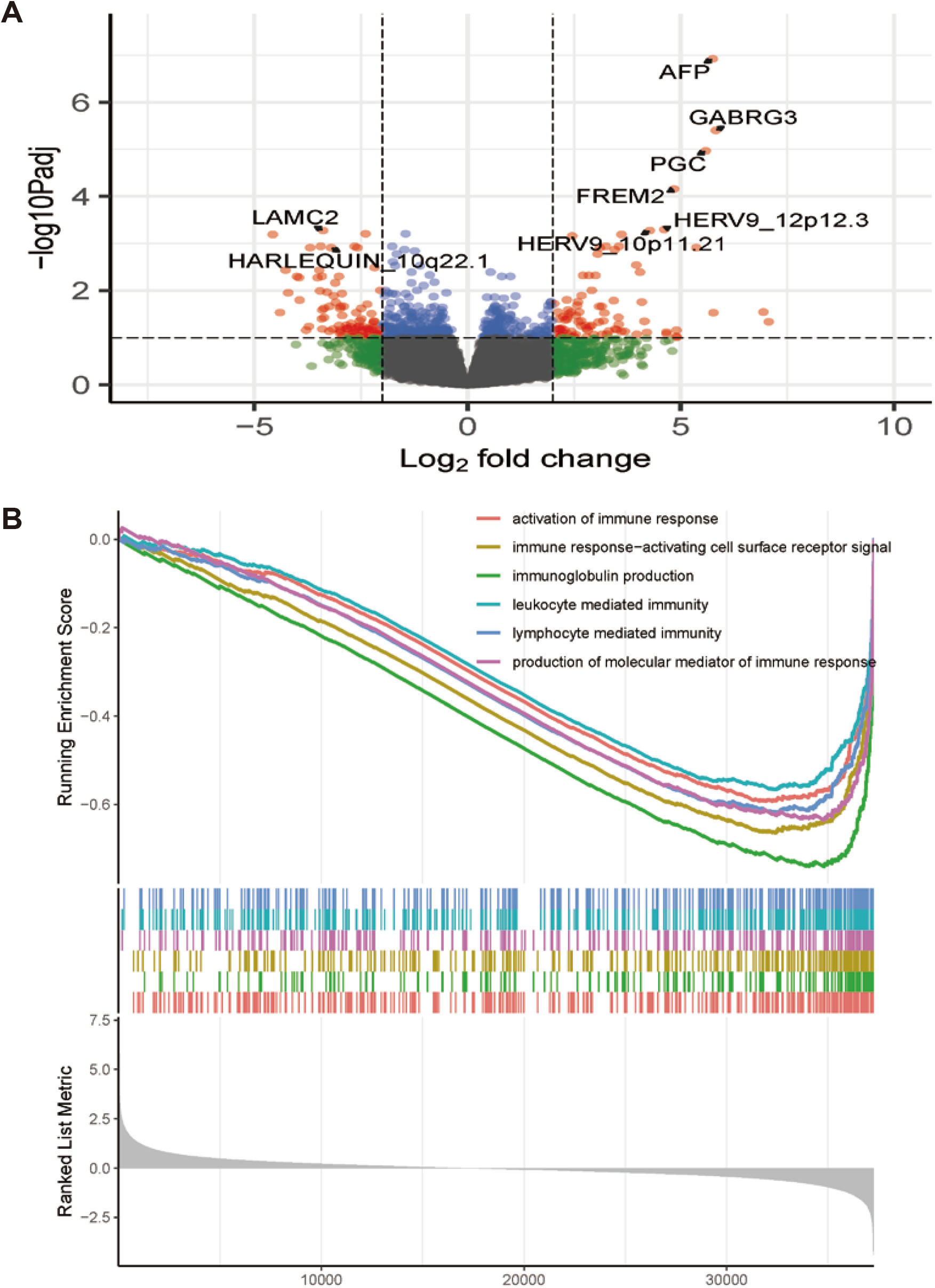
HBV- VS HBV+ HCC tumor.

We also investigated the differentially expressed genes between the triple negative (HBV, HCV, HDV negative) hepatocellular carcinoma and HBV positive (HCV, HDV negative) hepatocellular carcinoma. There are several human Endogenous Retrovirus are significantly up-regulated: ERVLE.13q21.1d 22.6 fold, HERVL.3q13.2 21.2 fold, HERVL.5p14.1a 20.6 fold and HERVLB4.4p12c 20.5 fold, as showed in Fig4A. GSEA showed activation of immune response, alpha/beta T cell activation, antigen receptor mediated signaling pathway, B cell activation pathways are dramatically downregulated in HBV positive hepatocellular carcinoma comparing to trip negative (HBV, HCV and HDV negative) hepatocellular carcinoma as showed in Fig4B.

**Figure 4:**
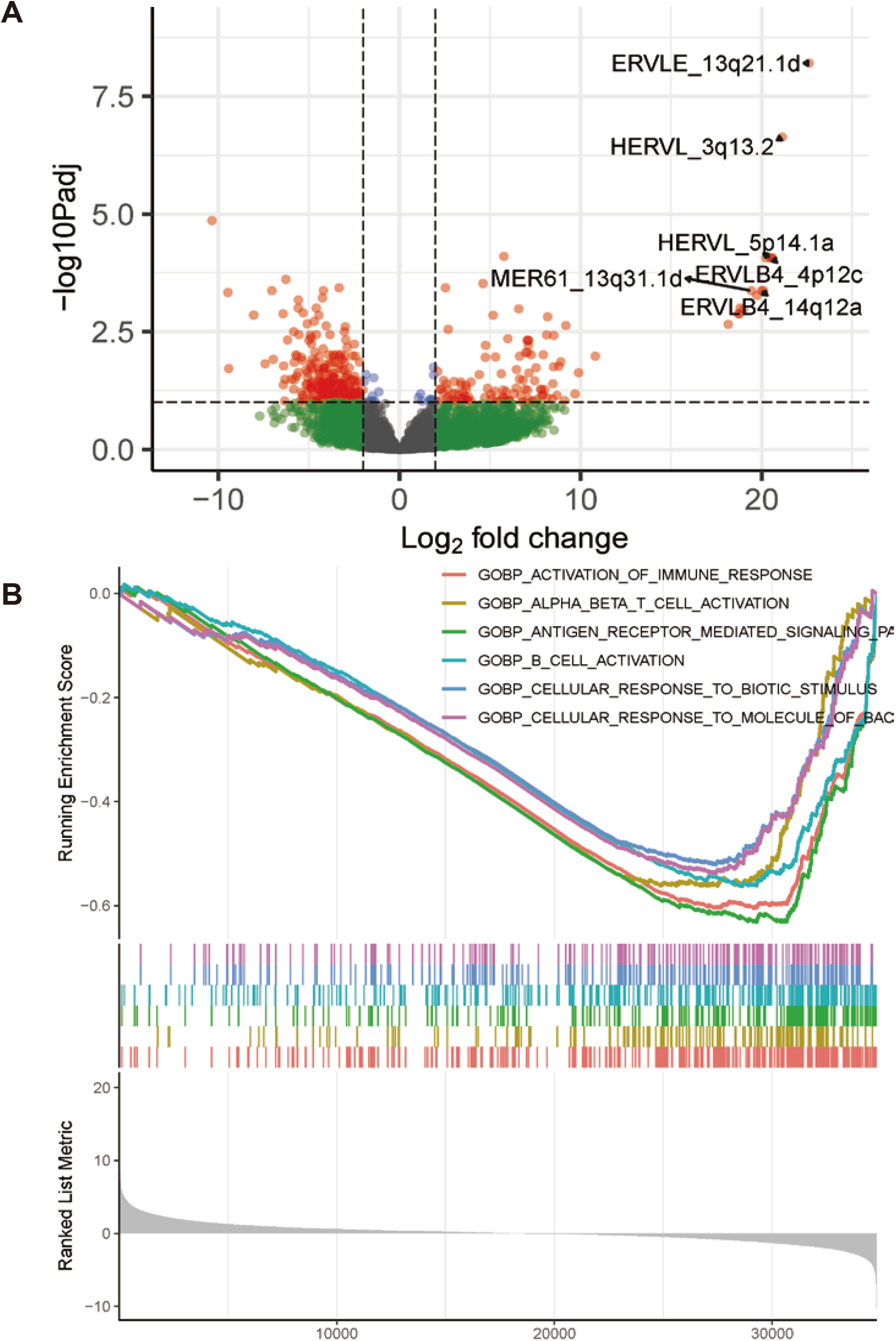
Trip Negative VS HBV+ (HCV-, HDV-) HCC tumor.

## Discussion

Single-cell RNA-seq (scRNA-seq) provides a cutting-edge method to study the cellular heterogeneity of the tumor microenvironment of many cancer types, by profiling the transcriptomics of thousands of individual cells. Bulk-tissue-level resolution may mask the complexity of alterations across cells and within cell groups, especially for less abundant cell types. ScRNA-seq analysis can help better understand cellular collective behavior and mutual regulatory mechanisms within a tissue ecosystem. Currently, studies on the relationship between hepatitis viruses and hepatocellular carcinoma at the single-cell level are still unclear. In our study, transcriptional level sequencing data of more than 70,000 single cells were collected and various cell types were analyzed, providing a new perspective for understanding the cell composition characteristics and pathogenesis development in the microenvironment of liver cancer. The microenvironment of cirrhotic and cancerized liver tissues are characterized with extensive immune infiltration. T cells and B cells are the most abundant and best-characterized population in tumor microenvironment (TME) of solid tumors. T cells are always the focus of anti-tumor activity research of HCC. A recent study found 11 T cell subsets of hepatocellular carcinoma based on their molecular characteristics through large-scale single-cell transcriptome sequencing. Meanwhile, the optimal efficacy of chimeric antigen receptor T cells in the immunotherapy of solid tumors inspired the research of CD8+T cells in hepatocellular carcinoma.

Human endogenous retroviruses (HERVs) are inherited genetic germline elements derived from exogenous retroviral infections throughout the evolution of the human genome, and account for 8 percent of our genome. The majority of Human endogenous retroviruses are defective due to evolutionarily acquired disruption or silencing mutations. Hence, no infectious activity remains from such Human endogenous retroviruses, but they may still be recognized as viral components by our immune system. Interestingly, elevated Human endogenous retroviruses expression has been associated with a variety of autoimmune disorders and cancers, although the causative roles and pathogenicity of Human endogenous retroviruses have not been clarified. Recent studies have shown that Human endogenous retroviruses expression is associated with tumor immune cytolytic activity12 as well as improved clinical response to immunotherapy in patients with cancers. Our results indicate that the malignant cells of certain myeloid origins harbor increased levels of certain Human endogenous retroviruses transcripts and could serve as a potential antigen pool for tumor-targeting immunotherapies. Furthermore, our data suggest that T cell recognition of Human endogenous retroviruses may support the immune reconstitution following HMA treatment, potentially by mediating cancer-cell control.

In this study, we discovered several Human endogenous retroviruses that can be potentially severed as biomarkers to indicating the hepatitis virus status in hepatocellular carcinoma. Also we found that the tumor microenvironment of hepatocellular carcinoma are affected by the hepatitis virus status too. These findings may shed light in the field of hepatitis virus and hepatocellular carcinoma.

## Supporting Information

If you intend to keep supporting files separately you can do so and just provide figure captions here. Optionally make clicky links to the online file using \href{url}{description}.

## Acknowledgements

We thank just about everybody.

## References

1. S. Anders, P. T. Pyl, and W. Huber. HTSeq–a Python framework to work with high-throughput sequencing data. Bioinformatics, 31(2):166–169, Jan 2015.

2. A. M. Bolger, M. Lohse, and B. Usadel. Trimmomatic: a flexible trimmer for Illumina sequence data. Bioinformatics, 30(15):2114–2120, Aug 2014.

3. J. Candia, E. Bayarsaikhan, M. Tandon, A. Budhu, M. Forgues, L. O. Tovuu, U. Tudev, J. Lack, A. Chao, J. Chinburen, and X. W. Wang. The genomic landscape of Mongolian hepatocellular carcinoma. Nat Commun, 11(1):4383, 09 2020.

4. C. D. Conde, C. Xu, L. B. Jarvis, D. B. Rainbow, S. B. Wells, T. Gomes, S. K. Howlett, O. Suchanek, K. Polanski, H. W. King, L. Mamanova, N. Huang, P. A. Szabo, L. Richardson, L. Bolt, E. S. Fasouli, K. T. Mahbubani, M. Prete, L. Tuck, N. Richoz, Z. K. Tuong, L. Campos, H. S. Mousa, E. J. Needham, S. Pritchard, T. Li, R. Elmentaite, J. Park, E. Rahmani, D. Chen, D. K. Menon, O. A. Bayraktar, L. K. James, K. B. Meyer, N. Yosef, M. R. Clatworthy, P. A. Sims, D. L. Farber, K. Saeb-Parsy, J. L. Jones, and S. A. Teichmann. Cross-tissue immune cell analysis reveals tissue-specific features in humans. Science, 376(6594):eabl5197, 2022.

5. A. Dobin, C. A. Davis, F. Schlesinger, J. Drenkow, C. Zaleski, S. Jha, P. Batut, M. Chaisson, and T. R. Gingeras. STAR: ultrafast universal RNA-seq aligner. Bioinformatics, 29(1):15–21, Jan 2013.

6. N. Grandi and E. Tramontano. Human Endogenous Retroviruses Are Ancient Acquired Elements Still Shaping Innate Immune Responses. Front Immunol, 9:2039, 2018.

7. Y. Hao, S. Hao E. Andersen-Nissen, W. M. M. III, S. Zheng, A. Butler, M. J. Lee, A. J. Wilk, C. Darby, M. Zagar, P. Hoffman, M. Stoeckius, E. Papalexi, E. P. Mimitou, J. Jain, A. Srivastava, T. Stuart, L. B. Fleming, B. Yeung, A. J. Rogers, J. M. McElrath, C. A. Blish, R. Gottardo, P. Smibert, and R. Satija. Integrated analysis of multimodal single-cell data. Cell, 2021.

8. C. Li, J. Chen, Y. Li, B. Wu, Z. Ye, X. Tian, Y. Wei, Z. Hao, Y. Pan, H. Zhou, K. Yang, Z. Fu, J. Xu, and Y. Lu. 6-Phosphogluconolactonase Promotes Hepatocellular Carcinogenesis by Activating Pentose Phosphate Pathway. Front Cell Dev Biol, 9:753196, 2021.

9. M. I. Love, W. Huber, and S. Anders. Moderated estimation of fold change and dispersion for RNA-seq data with DESeq2. Genome Biol, 15(12):550, 2014.

10. N. S. Mohammad, R. Nazli, H. Zafar, and S. Fatima. Effects of lipid based Multiple Micronutrients Supplement on the birth outcome of underweight pre-eclamptic women: A randomized clinical trial. Pak J Med Sci, 38(1):219–226, 2022.

11. M. Tarailo-Graovac and N. Chen. Using RepeatMasker to identify repetitive elements in genomic sequences. Curr Protoc Bioinformatics, Chapter 4:Unit 4.10, Mar 2009.

12. X. Wang, J. Park, K. Susztak, N. R. Zhang, and M. Li. Bulk tissue cell type deconvolution with multi-subject single-cell expression reference. Nat Commun, 10(1):380, 01 2019.

13. S. W. Wingett and S. Andrews. FastQ Screen: A tool for multi-genome mapping and quality control. F1000Res, 7:1338, 2018.

14. F. A. Wolf, P. Angerer, and F. J. Theis. SCANPY: large-scale single-cell gene expression data analysis. Genome Biol, 19(1):15, 02 2018.

15. T. Wu, E. Hu, S. Xu, M. Chen, P. Guo, Z. Dai, T. Feng, L. Zhou, W. Tang, L. Zhan, X. Fu, S. Liu, X. Bo, and G. Yu. clusterprofiler 4.0: A universal enrichment tool for interpreting omics data. The Innovation, 2(3):100141, 2021.

